# A novel sialylation site on *Neisseria gonorrhoeae* lipooligosaccharide links heptose II lactose expression with pathogenicity

**DOI:** 10.1101/302968

**Authors:** Sanjay Ram, Sunita Gulati, Lisa A. Lewis, Srinjoy Chakraborti, Bo Zheng, Rosane B. DeOliveira, George W. Reed, Andrew D. Cox, Jianjun Li, Frank St. Michael, Jacek Stupak, Xia-Hong Su, Sudeshna Saha, Corinna S. Landig, Ajit Varki, Peter A. Rice

**Author notes:** Corresponding author: Sanjay Ram, Division of Infectious Diseases and Immunology, University of Massachusetts Medical School, Lazare Research Building, Room 322, 364 Plantation Street, Worcester MA 01605, USA.

## Abstract

Sialylation of lacto-W-neotetraose (LNnT) extending from heptose I (HepI) of gonococcal lipooligosaccharide (LOS) contributes to pathogenesis. Previously, gonococcal LOS sialyltransterase (Lst) was shown to sialylate LOS in Triton X-100 extracts of strain 15253, which expresses lactose from both HepI and HepII, the minimal structure required for mAb 2C7 binding. Ongoing work has shown that growth of 15253 in cytidine monophospho-W-acetylneuraminic acid (CMP-Neu5Ac)-containing media enables binding to CD33/Siglec-3, a cell surface receptor that binds sialic acid, suggesting that lactose termini on LOS of intact gonococci can be sialylated. Neu5Ac was detected on LOSs of strains 15253 and a MS11 mutant with only lactose from HepI and HepII by mass spectrometry; deleting HepII lactose rendered Neu5Ac undetectable. Resistance of HepII lactose Neu5Ac to desialylation by α2-3-specific neuraminidase suggested an α2-6-linkage. Although not associated with increased factor H binding, HepII lactose sialylation inhibited complement C3 deposition on gonococci. 15253 mutants that lacked Lst or HepII lactose were significantly attenuated in mice, confirming the importance of HepII Neu5Ac in virulence. All 75 minimally passaged clinical isolates from Nanjing, China, expressed HepII lactose, evidenced by reactivity with mAb 2C7; mAb 2C7 was bactericidal against the first 62 (of 75) isolates that had been collected sequentially and were sialylated before testing. mAb 2C7 effectively attenuated 15253 vaginal colonization in mice. In conclusion, this novel sialylation site could explain the ubiquity of gonococcal HepII lactose *in vivo*. Our findings reiterate the candidacy of the 2C7 epitope as a vaccine antigen and mAb 2C7 as an immunotherapeutic antibody.

## Introduction

Gonorrhea affects about 78 million people annually worldwide (1); almost 470,000 of these are reported in the U.S. Multidrug-resistant gonorrhea has been reported on every continent and is a serious public health concern (2–6). Understanding how gonococci evade host immune defenses is an important step toward development of urgently needed safe and effective vaccines and novel therapeutics against this infection.

*N. gonorrhoeae* is a uniquely human-adapted pathogen (7). On a molar basis, lipooligosaccharide (LOS) is the most abundant gonococcal outer membrane molecule and plays a key role in many facets of pathogenesis (8–19). Host-like glycans expressed by lipooligosaccharide (LOS) of *N. gonorrhoeae* constitutes an example of molecular mimicry (8, 20).Two structures expressed by Neisserial LOS from heptose (Hep) I that mimic host glycans include lacto-N-neotetraose (LNnT; Galβ1-4GlcNAcβ1-3Galβ1-4Glcβ1-), identical to the terminal tetrasaccharide of paragloboside, a precursor of the major human blood group antigens (21), and globotriose (Galα1-4Galβ1-4Glcβ1-) that is identical to terminal globotriose trisaccharide of the P^*K*^-like blood group antigen (22). The seminal work of Harry Smith and colleagues showed that *N. gonorrhoeae* scavenge cytidine monophospho-*N*-acetylneuraminic acid (CMP-Neu5Ac) from its host to sialylate its LOS (23, 24). Both, LNnT and P^*K*^-like LOSs can be sialylated (25, 26). LOS sialylation inhibits complement activation and converts strains that are otherwise sensitive to killing by complement in serum to a ‘serum (or complement)-resistant’ phenotype (16, 27–29). Several other microbes also use sialic acid expression to their advantage to subvert host immunity (30–41), by mimicking host sialic acid-based “selfassociated molecular patterns”

Among members of the species Neisseria, the gonococcus uniquely expresses lactose extending from HepII. A monoclonal antibody (mAb) called 2C7 binds an epitope on LOS that requires expression of HepI and HepII lactose. Despite being undei control of a phase variable LOS glycosyltransferase *(lgt)* gene called *lgtG*, HepII lactose is expressed by ~95% of clinical isolates of *N. gonorrhoeae* (42), which suggests a key role in virulence. Isogenic mutant strains that lack *lgtG* show decreased virulence in the mouse vaginal colonization model of gonorrhea (43). Why HepII lactose promotes gonococcal virulence remains unclear. In recent work, we noted that growth of a gonococcal strain called 15253 that expresses lactose simultaneously from HepI and HepII (44) was capable of binding to Siglec-3 when grown in CMP-Neu5Ac-containing media (45). Siglec-3 binds exclusively to sialyoglycans (46, 47). These data suggested that lactose expressed by *N. gonorrhoeae* LOS could also be sialylated. This study describes sialylation of gonococcal LOS lactose termini and elucidates its function in complement evasion and virulence.

## Results

### Neu5Ac ‘caps’ *N. gonorrhoeae* HepII lactose

LOS glycan extensions from HepI and HepII for the strains used in this study are shown in Fig. 1. The ability of strains 15253 and MS11 2-Hex/G+, which express lactose extending from HepI and HepII, and their respective isogenic mutants, 15253/G– and MS11 2-Hex/G-, which express only lactose only from HepI, to add Neu5Ac to LOS was determined by SDS-PAGE. MS11 4-Hex/G-(expresses LNnT LOS from HepI) was used as a positive control for sialylation. An *lst* deletion mutant of 15253 (15253 *Δlst)* that lacks the ability to sialylate its LOS was also tested to address whether the previously described Lst enzyme is also responsible for sialylation of lactose.

**Fig. 1.**
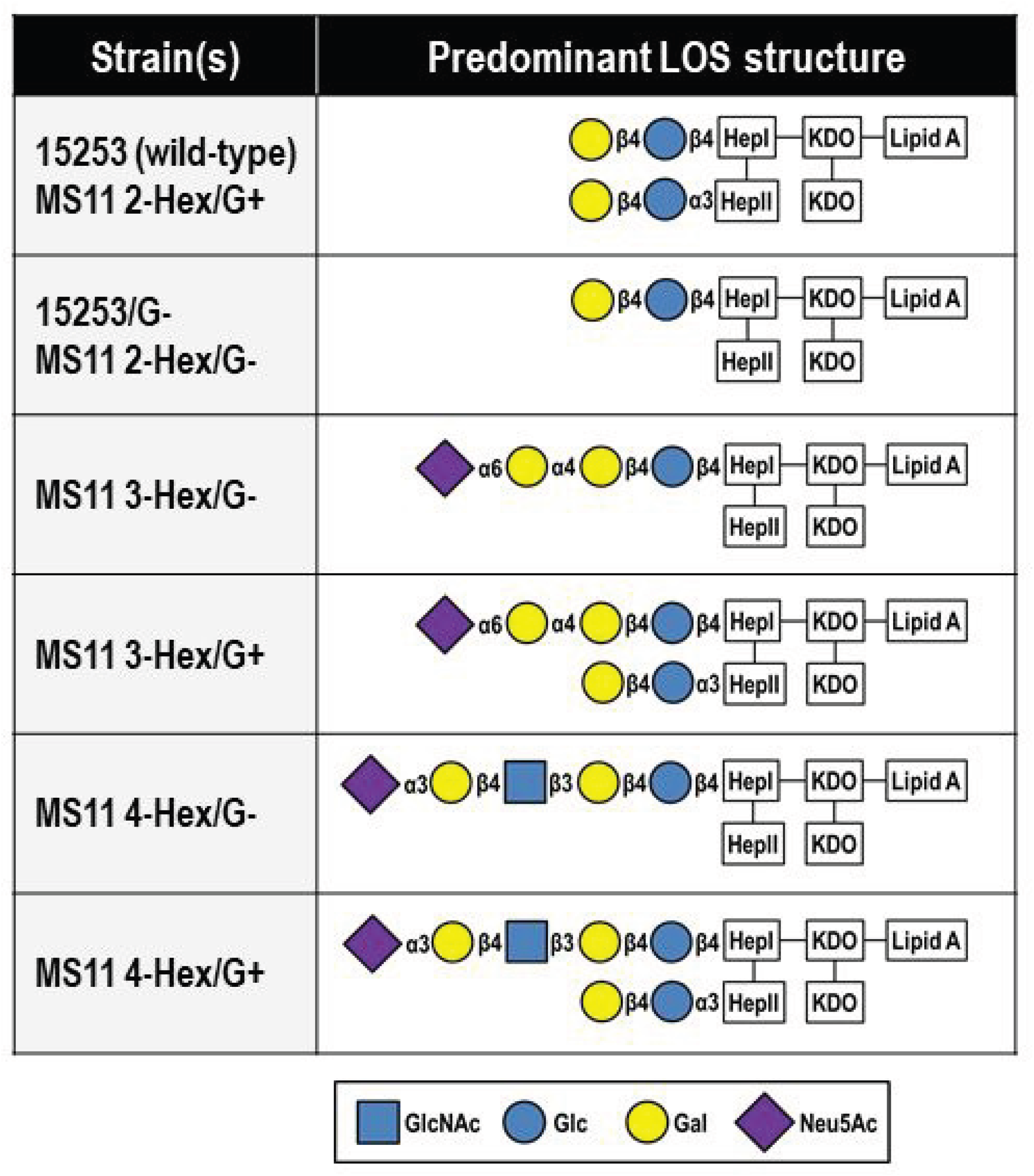
LOS glycan extensions from heptose (Hep) I and HepII elaborated by *N. gonorrhoeae* strains used in this study. Glycan extensions from HepI and HepII of the major LOS structure represented by the strains is shown using the symbol nomenclature for graphical representation of individual glycans (79). *N. gonorrhoeae* lack the ability to produce CMP-Neu5Ac, therefore capping of LOS with Neu5Ac requires the addition of CMP-Neu5Ac to growth media.

Bacteria were grown in media alone, or media supplemented with 100 μg/ml CMP-Neu5Ac for maximal LOS sialylation. As shown in Fig. 2, the upper band in the ‘+’ lane of 2-Hex/G+ shows slower mobility compared to the upper band in the corresponding ‘-’ lane. Note that despite fixing *lgtG* ‘on’, MS11 2-Hex/G+ expresses an LOS species with only HepI lactose (i.e., the LOS expressed by MS11 2-Hex/G-). This is because of export of LOS to the outer membrane prior to addition of the proximal Glc on HepII, as noted previously (48). Similarly, retarded mobility of 15253 LOS was also observed when grown in media containing CMP-Neu5Ac. The LOS of 4-Hex/G-(positive control for sialylation) incorporated Neu5Ac and migrated slower. There was no appreciable alteration in LOS migration when 2-Hex/G-, 15253/G– or 15253 *Δlst* were grown in CMP-Neu5Ac, suggesting that Neu5Ac was added to the terminal Gal of HepII lactose and that Lst was the enzyme responsible for sialylation.

**Fig. 2.**
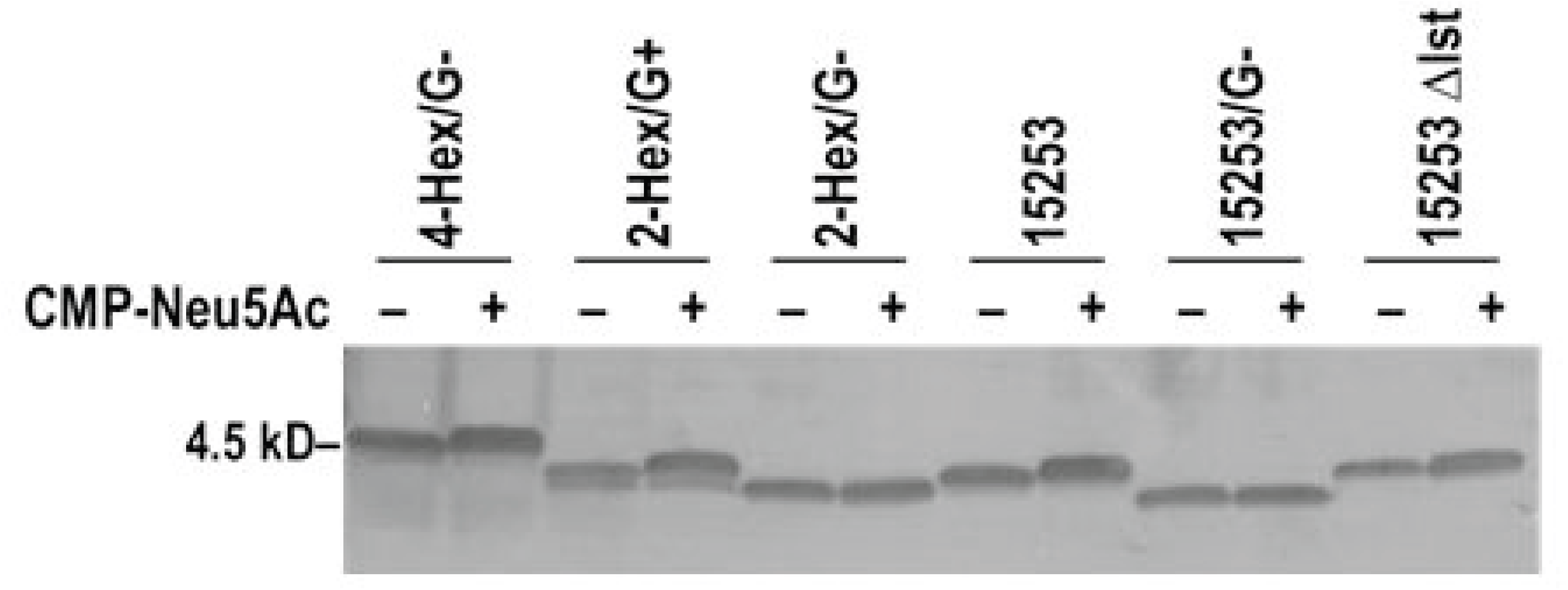
Evidence of sialylation of HepII lactose on *N. gonorrhoeae* 15253 and MS11 2-Hex/G+. *N. gonorrhoeae* 15253 and MS11 2-Hex/G+ both express lactose from HepI and HepII. Their mutants that lack HepII lactose were constructed by deleting *lgtG* (15253/G– and MS11 2-Hex/G-). 15253 Δlst lacks LOS sialyltransferase and cannot add Neu5Ac to LOS. MS11 4-Hex/G-expresses the sialylatable LNnT structure from HepI and served as a positive control for sialylation. All strains were in media with (+) or without (-) added CMP-Neu5Ac (100 μg/ml) for 2 h at 37 °C. Bacterial lysates were digested with protease K, separated on a 16% Tricine gel and LOS was visualized by silver staining. Retardation of LOS mobility following growth in CMP-Neu5Ac-containing media relative to LOS from bacteria grown in media devoid of CMP-Neu5Ac indicates sialylation.

Mass spectroscopic analysis of LOS purified from strains 15253, 15253/G-, MS11 2-Hex/G+ and MS11 2-Hex/G-grown in CMP-Neu5Ac and unsialylated 15253 (negative control) is shown in Supplemental Table S1. The data confirm the presence of sialic acid on 15253 and MS11 2-Hex/G+, but not on their isogenic mutants lacking HepII lactose. Collectively, the data strongly suggest that Neu5Ac is added to HepII lactose.

### Sialylation of HepII lactose does not enhance FH binding

Previously, we showed that sialylation of *N. gonorrhoeae* LNnT LOS, but not P^*K*^-like LOS, enhances human FH binding (26). We used strain MS11 to determine whether sialylation of HepII lactose enhances FH binding because this strain expresses PorB.1B and binds FH relatively weakly in the unsialylated state (49), which would more readily reveal increased FH binding with sialylation, if it were to occur. By contrast, strain 15253 (PorB.1A) binds high levels of FH even when unsialylated (49), which would limit the ability to detect an increase in FH binding with sialylation. FH binding to MS11 LOS mutants that expressed 2, 3 or 4 hexoses from HepI, each with (G+) or without (G-) HepII lactose was examined. The 3-Hex (P^*K*^-like LOS) and 4-Hex (LNnT) mutants served as negative and positive controls for FH binding with sialylation, respectively. As shown in Fig. 3, growth of the 2-Hex mutants in CMP-Neu5Ac did not enhance FH binding. Thus, enhanced FH binding to *N. gonorrhoeae* is restricted to LNnT LOS.

**Fig. 3.**
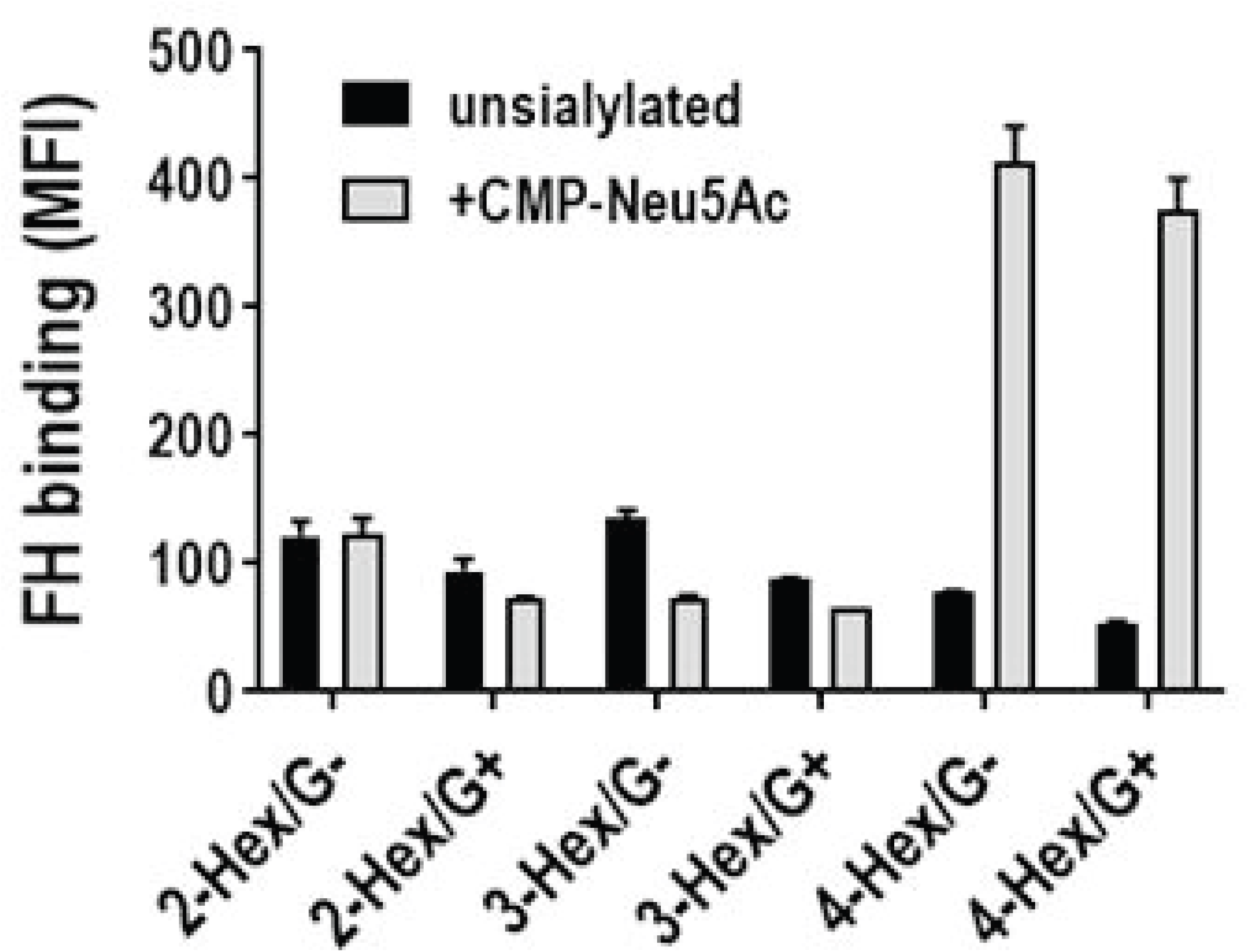
Enhanced FH binding upon LOS sialylation is restricted to strains that express the LNnT LOS structure. FH binding to isogenic LOS mutants of MS11 that express 2-Hex (lactose), 3-Hex (PK-like) or 4-Hex (LNnT) structures from HepI, with (G+) or without (G-) lactose extensions from HepII were grown in media alone or media containing CMP-Neu5Ac (25 μg/ml). Bacteria were incubated with FH (10 μg/ml) and bound FH measured by flow cytometry. Black bars, unsialylated bacteria; grey bars, sialylated bacteria. Control reactions, where FH was excluded, showed a median fluorescence below 10. Y-axis, median fluorescence (mean [range] of 2 separate observations).

### HepII lactose sialylation regulate complement activation

Neu5Ac capping of gonococcal LNnT and P^*K*^-like LOS both inhibit complement. *N. gonorrhoeae* bind C4BP and FH in a human-specific manner (50, 51). Initial attempts at measuring human C3 fragment deposition by flow cytometry on the two unsialylated strains using normal human serum revealed levels too low to discern the effects of sialylation on C3 deposition. Therefore, we used mouse complement, whose C4BP and FH do not bind *N. gonorrhoeae*, to study the effects of LOS sialylation on C3 deposition. Consistent with the addition of Neu5Ac to HepII lactose, C3 deposition decreased only on wild-type 15253 grown in media containing CMP-Neu5Ac (Fig. 4). Neither 15253/G-nor 15253 *Δlst* inhibited C3 deposition when grown in CMP-Neu5Ac, consistent with the inability of these two mutant strains to sialylate their LOSs.

**Fig. 4.**
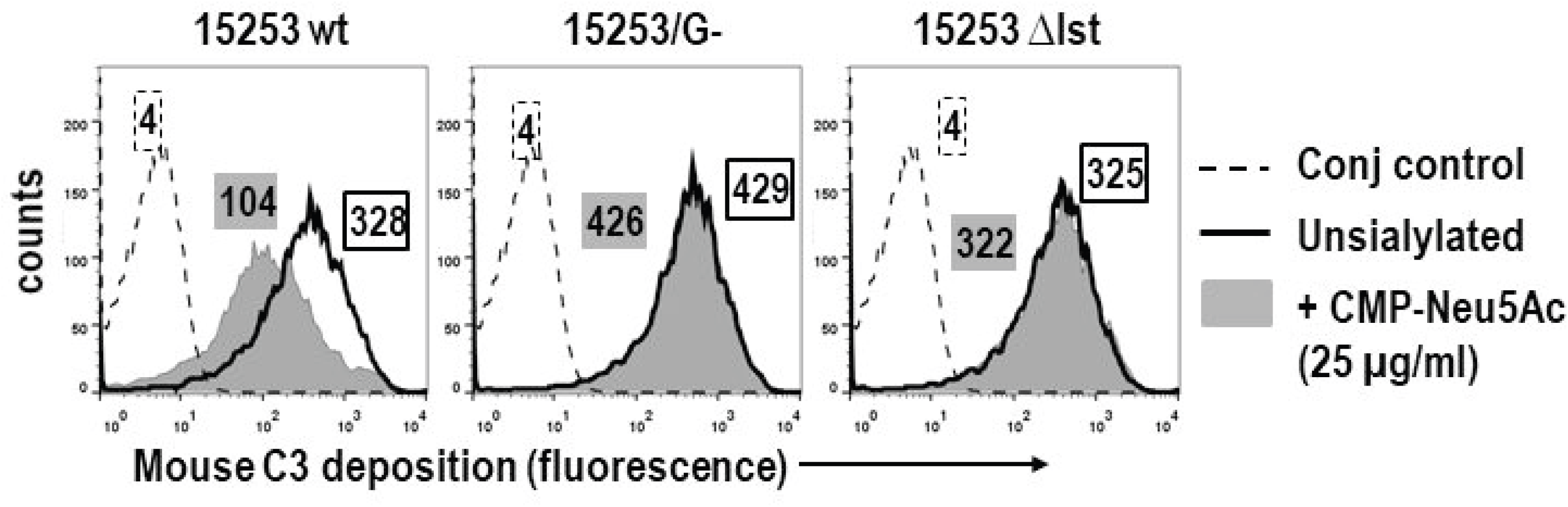
Sialylation of 15253 LOS inhibits complement activation. Strains 15253 and its isogenic mutant derivatives, 15253/G– (lacks HepII glycan extensions) and 15253 Δlst (lacks LOS sialyltransferase) were grown without or with CMP-Neu5Ac (25 μg/ml), incubated in 15% normal mouse serum for 20 min at 37 °C. C3 deposited on the bacterial surface was measured by flow cytometry. C3 deposited on bacteria grown in the presence or absence of CMP-Neu5Ac is shown by the grey shaded and solid black line histograms, respectively. Controls (no serum added) are shown by the broken lines. Numbers alongside histograms represent median fluorescence intensity (the border or shading of the text boxes that contain the numbers) correspond to that of the histograms). X-axis, fluorescence (log_10_ scale); Y-axis, counts. One representative experiment of at least two reproducible repeats is shown.

We next examined the effects of sialylation on complement activation on six MS11 LOS mutants (Fig. 5). The sialylatable 3-Hex and 4-Hex strains served as controls for Neu5Ac-mediated complement inhibition. Growth in media containing CMP-Neu5Ac decreased C3 deposition all tested strains. Similar to 15253, sialylation of MS11 2-Hex/G+ also inhibited C3 deposition. In contrast to 15253/G-, we noted a reproducible decrease in C3 deposition on the 2-Hex/G-mutant, suggesting that Neu5Ac was also added to the LOS of this strain. The degree of inhibition seen with sialylation of MS11 2-Hex/G– (a 4.1-fold decrease compared to the unsialylated parent, and 7.3-fold above baseline conjugate control levels) was less than that seen with complement inhibition upon sialylation of MS11 2-Hex/G+ (a 21-fold decrease compared to unsialylated 2-Hex/G+, and only 3-fold greater fluorescence compared to the baseline conjugate control). This amount of LOS sialylation of MS11 2-Hex/G-LOS, although functional, was too small to be appreciated by changes in mobility on a Tricine gel or by MS analysis.

**Fig. 5.**
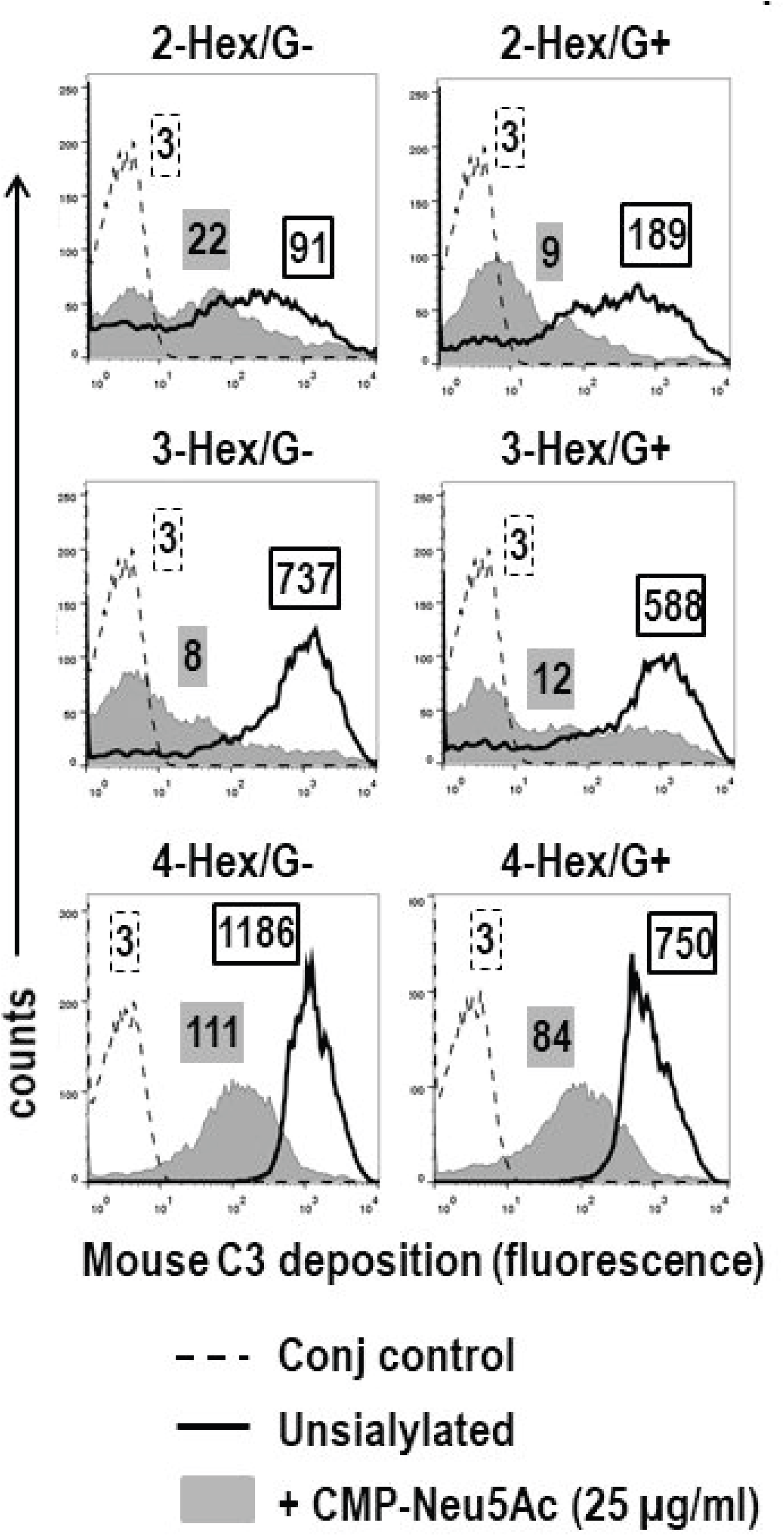
Complement inhibition by MS11 mutants that express lactose LOS extensions. Isogenic MS11 mutants that express predominantly lactose (2-Hex), PK structure (3-Hex) or LNnT (4-Hex) from HepI, with (G+) or without (G-) lactose from HepII, were grown in the absence or presence of CMP-Neu5Ac (25 μg/ml) and were incubated with 15% normal mouse serum for 20 min at 37 °C. Mouse C3 deposited on the bacterial surface was measured by flow cytometry. C3 deposited on bacteria grown in the presence or absence of CMP-Neu5Ac is shown by the grey shaded areas and solid black lines, respectively. Controls (no serum added) are shown by the broken lines. Numbers alongside histograms represent median fluorescence intensity (the border or shading of the text boxes that contain the numbers) correspond to that of the histograms). X-axis, fluorescence (log_10_ scale); Y-axis, counts. One representative experiment of at least two reproducible repeats is shown.

To discern the sialic acid linkage to HepII substituted lactose we examined the effect of α2-3-linkage-specific neuraminidase on mouse C3 deposition (Fig. 6). Sialylated 2-Hex/G+ and the corresponding wild-type strain 15253 that possessed the same pattern of Hep I and Hep II hexose substitutions, failed to show increased C3 deposition after treatment with recombinant α2-3-specific sialidase followed by incubation with 15% normal mouse serum; resistance to α2-3-sialidase suggests an α2-6 linkage.

**Fig. 6.**
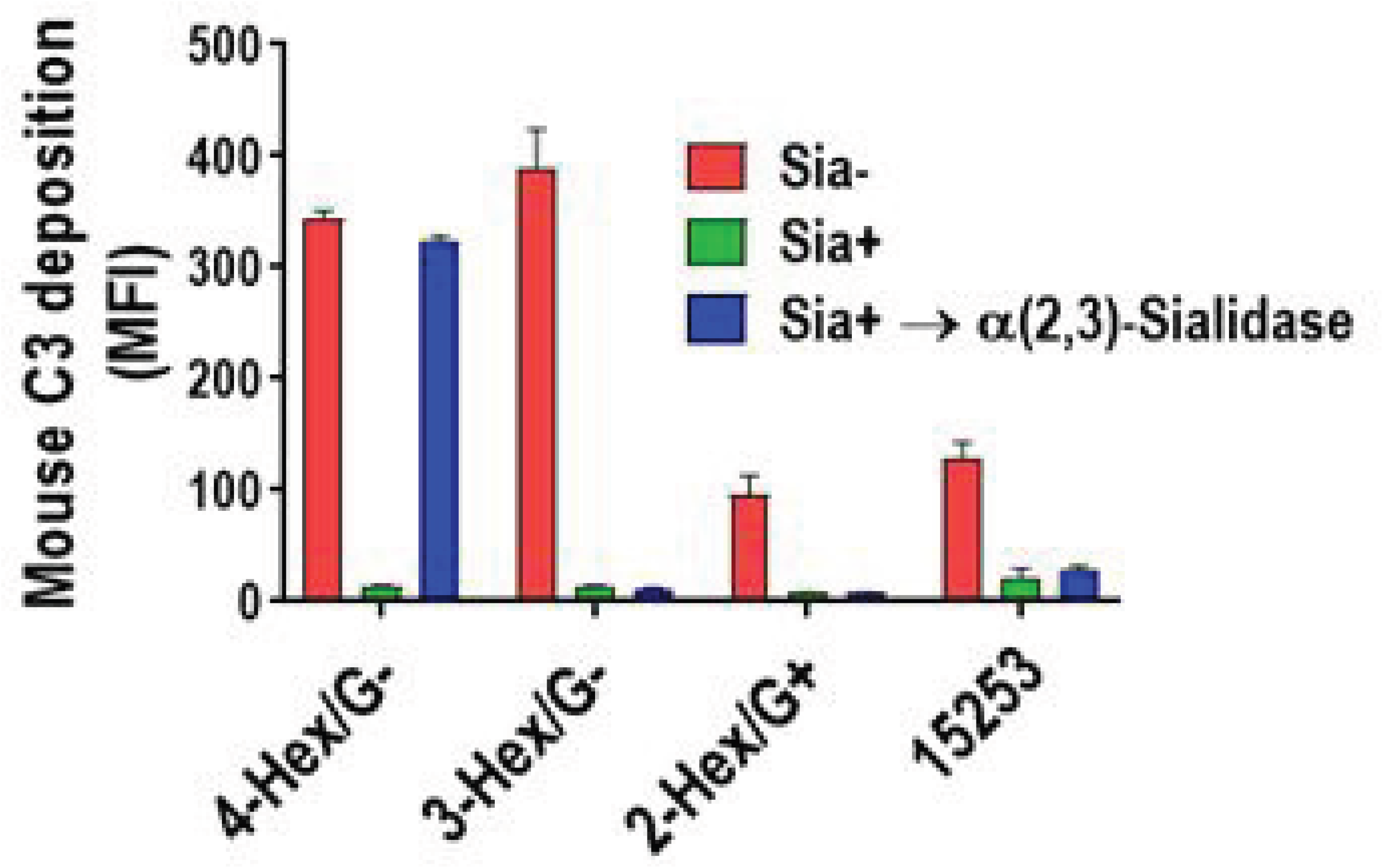
Neu5Ac added to HepII lactose resists removal by α2-3-sialidase. MS11 2-Hex-G+ and 15253 were grown in the absence or presence of CMP-Neu5Ac (25 μg/ml). MS11 4-Hex/G-, which expresses LNnT that is sialylated through an α2-3-linkage and MS11 3-Hex/G-, which expresses the P^*K*^-like LOS that becomes sialylated through an α2-6-linkage, were used as positive and negative controls for desialylation, respectively. Bacteria were treated with recombinant α2-3-specific sialidase or with neuraminidase (sialidase) buffer alone, then incubated with 15% normal mouse serum for 20 min at 37 °C. Mouse C3 deposited on bacteria (shown as median fluorescence intensity (MFI) on the Y-axis) was measured by flow cytometry. Controls (no added serum) showed fluorescence less than 10 units and have been omitted for simplicity. Each bar represents the mean (range) of two separate experiments.

### Loss of sialic acid on HepII lactose impairs *N. gonorrhoeae* vaginal colonization in mice

The ability of 15253 (wild-type), 15253/G– and 15253 Δlst to colonize the genital tract of *Cmah* knockout (KO) mice was compared. *Cmah* KO mice lack the enzyme CMP-N-acetylneuraminic acid hydroxylase (CMAH) and akin to humans, cannot convert Neu5Ac to Neu5Gc. Thus, these mice provide a ‘human-like’ sialic acid milieu to study the effects of LOS sialylation on virulence. *Cmah* KO mice support *N. gonorrhoeae* colonization slightly better than control wild-type BALB/c mice (52). Loss of either HepII lactose or Lst significantly attenuated the duration and burden of bacterial colonization (Fig. 7A-C). These data provide strong evidence for the importance of sialylation of HepII lactose in gonococcal virulence. We were unable to evaluate the effects of LOS sialylation in MS11 2-Hex/G+ because this strain colonized mice for only 3 days (data not shown).

**Fig. 7.**
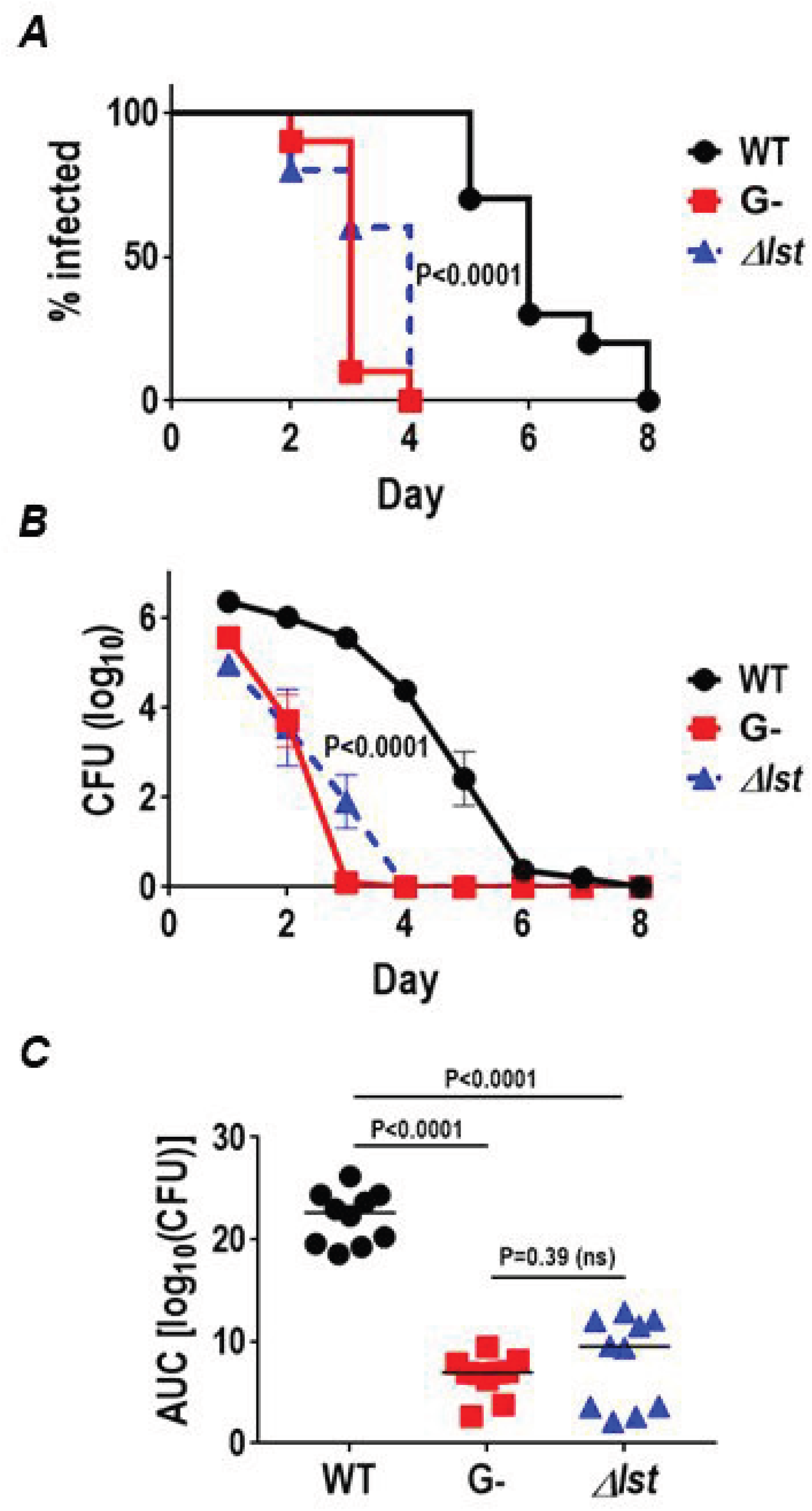
Sialyation of HepII lactose enhances virulence of strain 15253. Cmah knockout mice that express only Neu5Ac (the form of sialic acid found in humans), but not Neu5Gc (the form in wild-type mice), were infected with wild-type (WT) *N. gonorrhoeae* 15253 (5.5 × 10^7^ CFU) and its isogenic mutants 15253/G– (lacks any HepII glycan extension; 4.3 × 10^7^ CFU) and 15253 *Δlst* (lacks LOS sialyltransferase; 4.9 × 10^7^ CFU) (n=10 mice per group). Vaginas were swabbed daily to enumerate *N. gonorrhoeae* CFUs. ***A***. Kaplan Meier curves showing time to clearance. WT bacteria versus G– and WT versus *Δlst*, P<0.0001 by Mantel-Cox log-rank test. ***B***. CFU versus time. X-axis, day; Y-axis, CFU (log_10_). ***C***. Area Under Curve (AUC) analysis for consolidated bacterial burden over time. Pairwise comparisons between G– and *Δlst* with the control group were made by Mann-Whitney’s non-parametric t test. Comparisons across groups were made by one-way ANOVA (Kruskal-Wallis non-parametric test; P<0.0001).

### Effect of sialylation on mAb 2C7 binding and efficacy

mAb 2C7 targets a LOS epitope being developed as a gonococcal vaccine candidate. The minimal LOS structure required for mAb 2C7 binding are lactoses simultaneously extending from both HepI and HepII. Glycan extensions beyond lactose on HepII, for example with GalNAc-Gal seen in a mutant strain selected under pyocin pressure called JW31R, abrogates mAb 2C7 binding (53). We therefore asked whether sialylation of HepII lactose affected mAb 2C7 binding and function. While sialylation of 15253 did not affect mAb 2C7 binding (Fig. 8A, left graph), sialylation of MS11 2-Hex/G+ resulted in a reproducible ~2– to 3-fold reduction in mAb 2C7 binding (Fig. 8A, right graph). Similar binding of mAb 2C7 to sialylated and unsialylated 15253 allowed us to assess the functional effects of HepII lactose sialylation when antibody binding was kept constant. As shown in Fig. 8B, increasing amounts of CMP-Neu5Ac in media caused a dose-dependent decrease in killing by mAb 2C7.

**Fig. 8.**
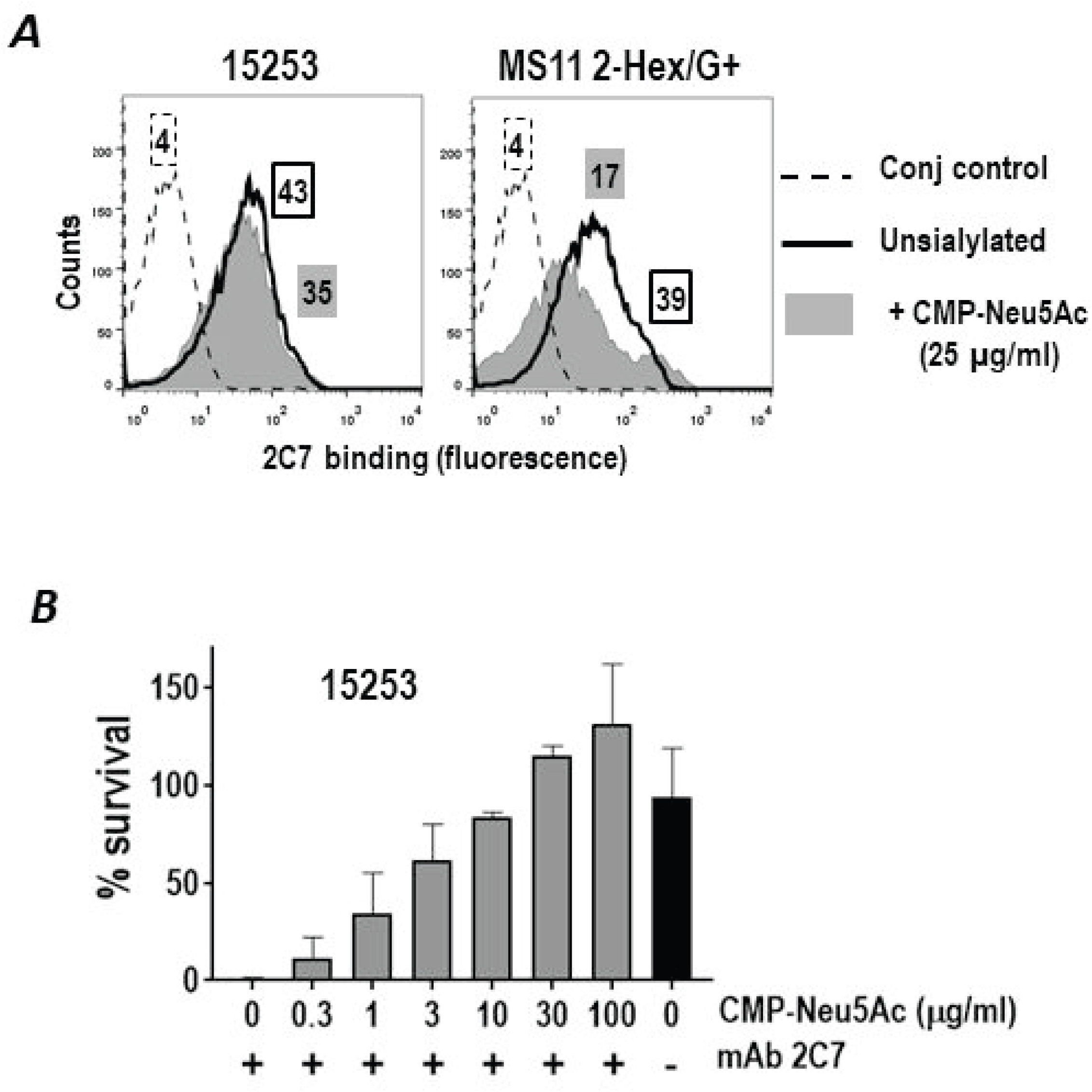
Effect of HepII lactose sialylation on the binding and bactericidal efficacy of mAb 2C7. A. Sialylation decreases binding of mAb 2C7 to MS11 2-Hex/G+, but not to 15253. *N. gonorrhoeae* were grown in media alone or media supplemented with 25 μg/ml CMP-Neu5Ac and binding of mAb 2C7 (10 μg/ml) to 15253 (left graph) and MS11 2-Hex/G+ was measured by flow cytometry. The solid black line shows mAb 2C7 binding to unsialylated bacteria; the grey shaded histogram, mAb 2C7 binding to sialylated bacteria. The control indicates bacteria incubated with anti-mouse IgG-FITC (no added mAb 2C7). One experiment of two reproducible repeats is shown. B. Addition of CMP-Neu5Ac to growth media in increasing concentrations decreases the bactericidal efficacy of mAb 2C7 against *N. gonorrhoeae* 15253. Serum bactericidal assays were performed with 20% pooled normal human serum (NHS) as the complement source. Where indicated, mAb 2C7 was added to a concentration of 10 μg/ml. Y-axis, percent survival following incubation of the reaction for 30 min relative to survival at 0 min.

### mAb 2C7 is active against strain 15253 in *vivo*

In light of prior work that showed the importance of LOS sialylation for infection of mice (54, 55) and the observed resistance of sialylated 15253 to mAb 2C7 *in vitro* (Fig. 8), we examined the efficacy of mAb 2C7 versus 15253 in the BALB/c mouse vaginal colonization model. A ‘passive immunization model’ to address the efficacy of mAb 2C7 to simulate effects of vaccine antibody was used (43). Wild-type BALB/c mice (n=8) were administered mAb 2C7 10 μg intraperitoneally on Days −2, −1 and 0, and CFUs were monitored daily. The control group (n=7) received mouse IgG3. mAb 2C7 significantly shortened the duration and burden of infection with 15253 (Fig. 9A-C).

**Fig. 9.**
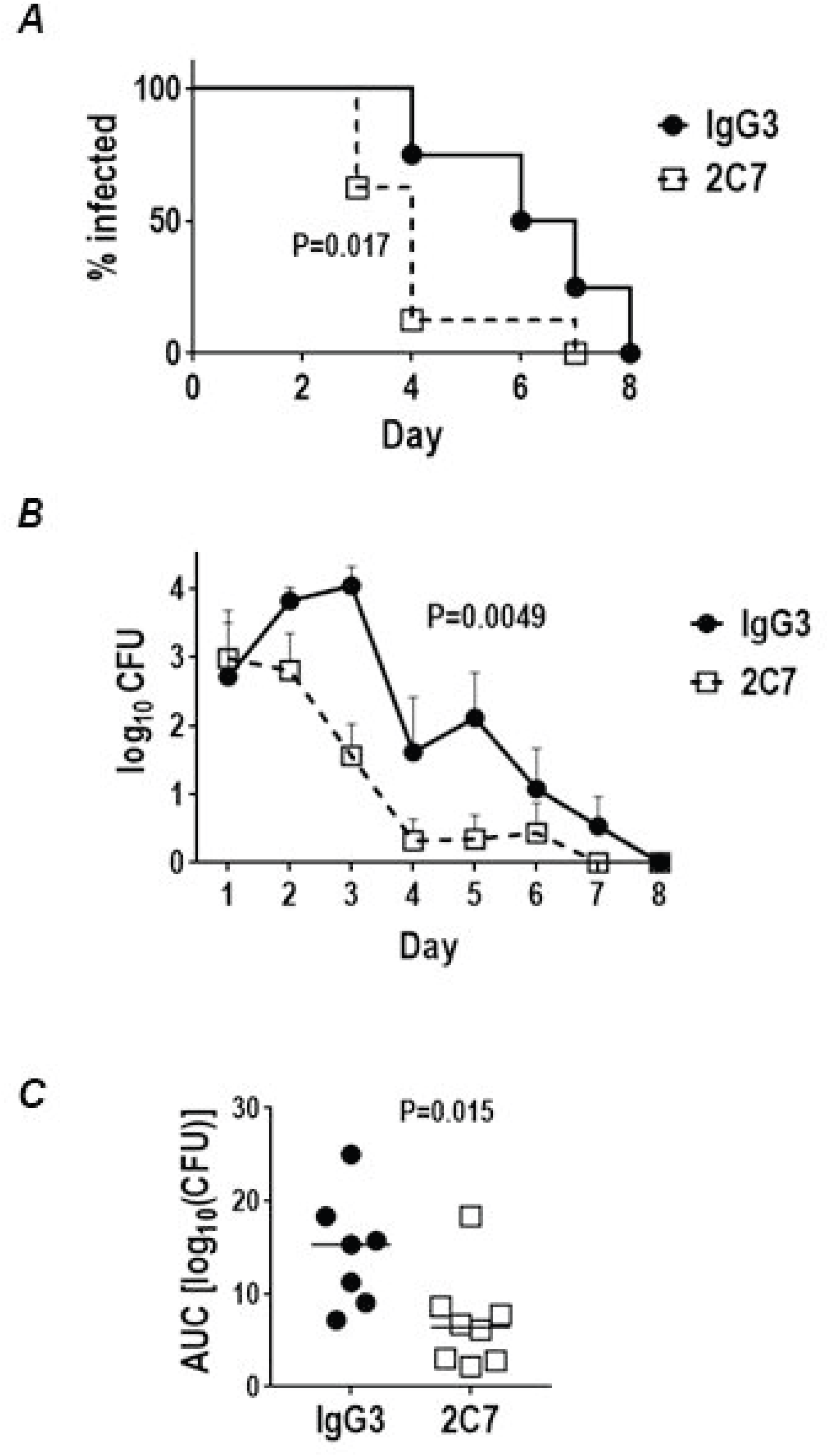
mAb 2C7 is attenuates infection with *N. gonorrhoeae* in the murine vaginal colonization model. Wild-type BALB/c mice were treated with either mAb 2C7 (10 μg intraperitoneally twice a day on days −2, −1 and 0) or a similar dose of control mouse IgG3 and then infected with 5 × 10^5^ CFU of 15253. Vaginas were swabbed daily to enumerate CFUs. A. Kaplan Meier curves showing time to clearance. The two groups were compared using the Mantel-Cox log-rank test. B. CFU versus time. X-axis, day; Y-axis, CFU (log_10_). C. Area Under Curve (AUC) analysis showing consolidated bacterial burdens over time. Pairwise comparisons between the two groups were made by Mann-Whitney’s non-parametric t test.

Expression of the 2C7 epitope by contemporary clinical isolates of *N. gonorrhoeae*. Despite being under control of a phase variable gene, *lgtG*, the 2C7 LOS epitope (Fig. 10A) was expressed by 94% of gonococci recovered directly from cervical secretions from a cohort of women who attended a sexually transmitted disease (STD) clinic in Boston (42). We examined a collection of minimally (≤3) passaged isolates cultured from the female contacts of men with gonorrhea who were referred to a STD clinic in Nanjing, China, for expression of the 2C7 LOS epitope by whole cell ELISA. We also examined isolates for their ability to bind to mAb L8 (recognizes an epitope defined by HepI lactose and phosphoethanolamine [PEA] substitution at the 3-position on HepII; expression of HepII lactose abrogates mAb L8 binding) and mAb 3F11 (recognizes terminal [unsialylated] lactosamine of LNnT) (Fig. 10A). As shown in Fig. 10B, each of 75 isolates bound to mAb 2C7, albeit to varying degrees, as did mAb L8 and 3F11. mAb L1 barely bound to any of the tested isolates.

**Fig. 10.**
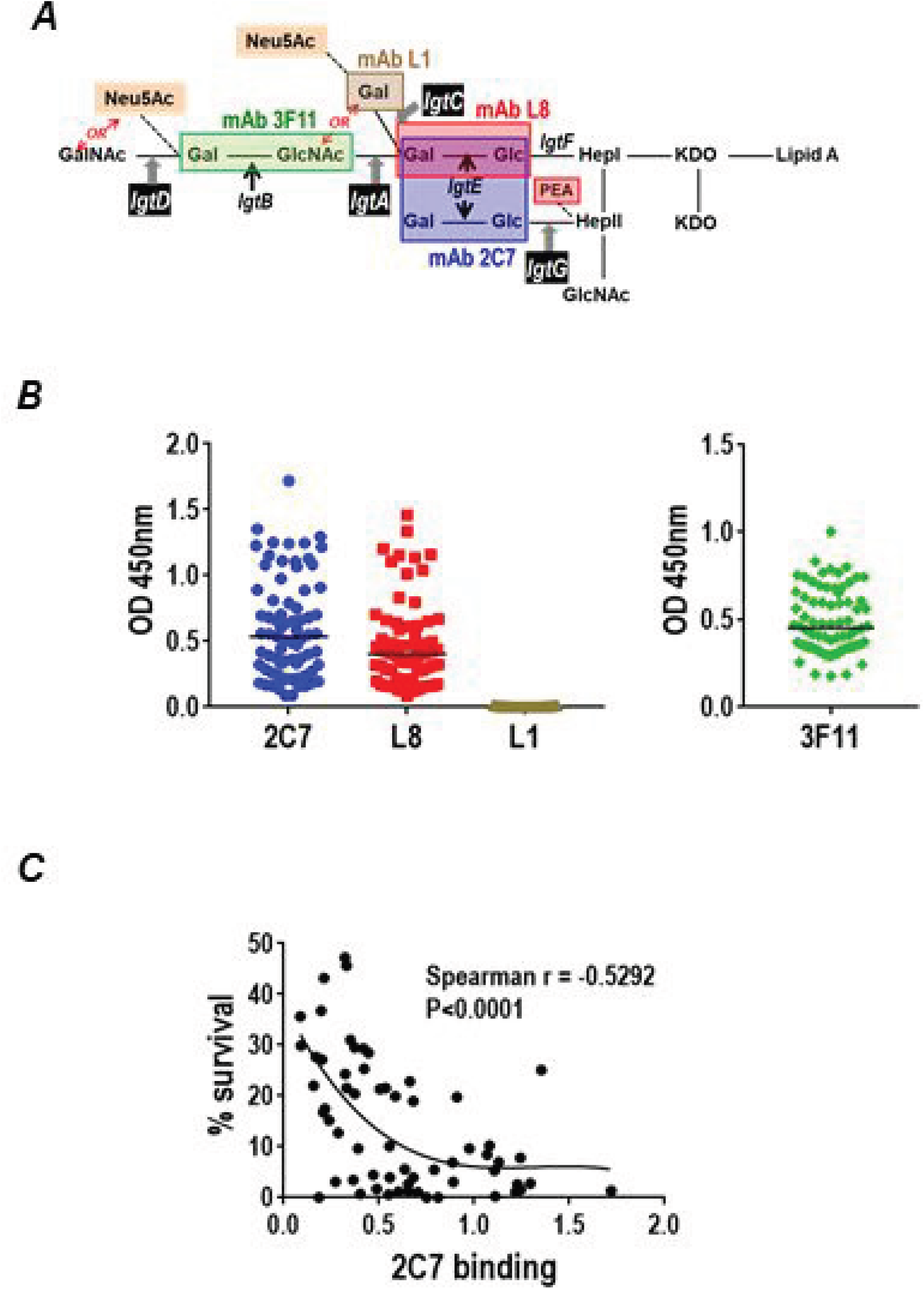
Expression of the 2C7 LOS epitope by clinical isolates from Nanjing, China and bactericidal efficacy of mAb 2C7. A. Schematic showing reactivity of anti-LOS mAbs 2C7, L1, L8 and 3F11. mAb 2C7 requires expression of lactose from HepI and HepII simultaneously (53). mAb L8 recognizes lactose from HepI in conjunction with a phosphoethanolamine (PEA) at the 3-position of HepII (74). Expression of 3-PEA from HepII requires *lgtG* to be phase-varied ‘OFF’, thus binding of mAb 2C7 and L8 occur exclusively and do not bind to overlapping epitopes. mAb 3F11 binds to unsialylated terminal lactosamine of the LNnT structure; any extension beyond lactosamine – for example with GalNAc *[lgtD* phase-varied ‘ON’] or the addition of Neu5Ac by adding CMP-Neu5Ac to growth media – abrogates mAb 3F11 binding (73). mAb L1 binds to the P^*K*^-like globotriose structure (Galα(1,4)-Galβ(1,4)-Glc) (75). B. Reactivity of mAbs 2C7, L1, L8 and 3F11 to 75 minimally passaged *N. gonorrhoeae* isolates recovered from men with urethritis attending the Nanjing (China) STD clinic. Binding of mAbs was determined by whole-cell ELISA. mAbs 2C7, L1 and L8 are all mouse IgG, while 3F11 is IgM, therefore shown as a separate graph. C. Complement-dependent bactericidal activity of mAb 2C7 against the first 62/75 isolates collected from men with urethritis in a Nanjing (China) study of gonococcal transmission from men to women as a function of mAb 2C7 binding. Bacteria were grown in media containing CMP-Neu5Ac (2 μg/ml) to enable them to fully resist killing (>100% survival) by 16.7% normal human serum (NHS). Survival of bacteria at 30 min following incubation with mAb 2C7 (5 μg/ml) plus NHS (16.7%) is shown as a function of mAb 2C7 binding (X-axis).

We next assessed the ability of mAb 2C7 to mediate complement-dependent bactericidal activity against the first 62 of 75 isolates collected from men with urethritis in a Nanjing (China) study of gonococcal transmission from men to women. Because some of the strains were sensitive to killing by 16.7% pooled normal human serum (NHS) that was used as the complement source, all isolates were grown in media containing 2 μg/ml CMP-Neu5Ac to render them fully serum resistant (>100% survival). All (100%) of isolates were killed >50% in the presence of 5 μg/ml of mAb 2C7 and NHS. This included two of four isolates that bound very low levels of mAb 2C7 by ELISA (OD_450nm_ between 0.065 and 0.090). Further, serum bactericidal activity correlated with levels of mAb 2C7 binding (Fig. 10C).

## Discussion

The novel finding in this report is the presence of Neu5Ac on *N. gonorrhoeae* HepII lactose. To our knowledge, *N. gonorrhoeae* is the only member of the genus Neisseria that expresses lactose extending from HepII. Certain *N. meningitidis* strains possess *lgtG* and can substitute Glc at the 3 position of HepII (seen in LOS immunotypes L2 and L4 (56, 57)), but extensions beyond the proximal Glc in meningococci have not been described. Prior work by Mandrell et al provided evidence for the ability of 15253 Lst to sialylate lactose, although not in context of intact bacteria; LOS in Triton X-100 extracts of strain 15253 (which also contains Lst) could incorporate radiolabeled Neu5Ac when supplied with exogenous CMP-[^14^C]-Neu5Ac (58).

mAb 2C7 recognized 94% of 68 gonococci examined directly from cervical secretions express and 95% of 101 randomly chosen fresh (second passage) gonococcal isolates from a sexually transmitted disease clinic in Boston (42). We recently surveyed 75 minimally passaged gonococcal isolates from Nanjing, China and noted that 100% of isolates reacted with mAb 2C7. All strains also expressed LNnT, which suggests that both sialylatable glycans are important for gonococcal pathogenesis. The importance of LNnT sialylation, both in humans and in the mouse vaginal colonization model has been established (12, 54, 55, 59). Phase variability of *IgtA* and *IgtD* control expression of LNnT (17). *IgtA* and *IgtC* both ‘off’ would result in expression of lactose, while the combination of *IgtA* off and *IgtC* on would result in elaboration of the P^*K*^-like (3-Hex) structure. If *IgtA* and *IgtD* are both ‘on’, GalNAc is added to the terminal Gal of LNnT and prevents sialylation. Sialic acid likely plays a multifaceted role in Neisserial pathogenesis. In addition to inhibiting complement, enhancing resistance to opsonophagocytosis and cationic antimicrobial peptides (60, 61), Neu5Ac engages sialic acid-binding immunoglobulin-type lectins (Siglecs) many of which are, in turn, linked to an immunoreceptor tyrosine-based inhibition (ITIM) motif and inhibit the inflammatory response (62). Neu5Ac has also been identified in gonococcal biofilms (63). Because it can also sialylate HepII lactose, the gonococcus has the capacity to maintain LOS sialylation even when the previously described sialylatable LNnT or P^*K*^-like structures is not expressed from HepI. While HepII lactose can be sialylated when HepI also expresses lactose, it is unclear whether HepII lactose can be sialylated when LNnT or P^*K*^ is also expressed on HepI. Gilbert and colleagues showed that meningococcal Lst could add Neu5Ac to 6(5-fluorescein-carboxamido)– hexanoic acid succimidyl ester (FCHASE)-aminophenyl-lactose. Lst added ~6.4-fold or ~3.2-fold more Neu5Ac onto lactosamine (LNnT is lactosamine-lactose) compared to lactose at substrate concentrations of 0.2 mM or 1.0 mM, respectively (64). Based on these data, we speculate that LNnT is preferentially sialylated over lactose when both glycan species are expressed.

The importance of Neu5Ac on HepII lactose in pathogenesis was illustrated by attenuation of 15253 Δlst in the mouse vaginal colonization model. Thus, unsialylated HepII lactose does not support virulence in this model. Expression of HepII lactose by almost all clinical isolates highlights the importance of maintaining *lgtG* ‘on’ *in vivo*. We are not aware of any naturally occurring gonococcal isolate that lacks *lgtG*. We have shown previously that a *lgtG* deletion mutant of *N. gonorrhoeae* FA1090 was less virulent than its wild-type parent (43). Lam and Gray-Owen showed that serial passage of *N. gonorrhoeae* in mice resulted an increased fraction of mice infected with each subpassage and in a reproducible selection of variants with lgtG ‘on’, providing further strong evidence of the importance of HepII lactose expression in vivo (65).

We noted that MS11 2-Hex/G-inhibited mouse complement when grown in CMP-Neu5Ac. The amount of Neu5Ac incorporation onto HepI lactose in this mutant was likely too small to be detected by shifts on SDS-PAGE or by MS analysis, but was nevertheless sufficient for functional activity, limited as it was. In contrast, 15253/G-, which also expresses only lactose from HepI, did not inhibit mouse C3 deposition when grown in CMP-Neu5Ac-containing media, suggesting that the extent and influence of HepI lactose sialylation on function may differ across strains. Whether a difference in HepI lactose sialylation exists between the two strains is unclear despite differences in function, but could relate to differences in Lst sequence and/or levels of Lst activity. Translation of DNA sequences of the *lst* ORF of 15253 and MS11 showed a single amino acid sequence difference; 15253 possessed a Q (seen in 16 other *N. gonorrhoeae* Lst sequences), while MS11 possessed an E (seen in >400 *N. gonorrhoeae* Lst sequences) at position 266. The −35/−10 promoter sequence, transcription start sites and the Shine-Dalgarno sequence were identical in 15253 and MS11. Packiam et al showed wide variation in *lst* mRNA levels across gonococcal strains, but mRNA levels often did not correlate with Lst activity as measured by sialylation of Triton X-100 bacterial extracts (66).

Linkage of Neu5Ac is a key determinant of its ability to interact with the C-terminus of FH. Blaum and colleagues showed that the interaction between sialic acid and FH domain 20 is restricted only to α2-3-linked Neu5Ac; α2-6– or α2-8-linked Neu5Ac do not interact with FH (67). The Neu5Ac-lactose bond is resistant to α2-3 specific sialidase, suggesting an α2-6 linkage. In accordance with the findings of Blaum et al, sialylation of HepII lactose – presumably through an α2-6 linkage – did not increase FH binding to *N. gonorrhoeae*. We acknowledge that further structural characterization is necessary to confirm the nature of the Neu5Ac-lactose linkage. How Neu5Ac on HepII lactose regulates complement remains unclear. Similar to LNnT sialylation, Neu5Ac linked to LNnT may also inhibit the classical pathway by reducing binding of IgG directed against select surface targets to the bacterial surface.

Despite similar amounts of mAb 2C7 binding to sialylated compared to unsialylated 15253, the sialylated derivative was resistant (>50% survival) to mAb 2C7 plus human complement when exposed to CMP-Neu5Ac concentrations ≥3 μg/ml. A possible explanation is that targets for C4b and C3b on LOS (68), may be obscured by the presence of Neu5Ac, thereby diminishing Ab efficacy. However, mAb 2C7 remained effective against 15253 in the mouse vaginal colonization model, where the organism is sialylated and additional factors such as opsonophagocytosis may contribute to its bactericidal activity. Mouse FH and C4BP do not bind to gonococci (50, 51). Therefore, the barrier that mAb 2C7 must surmount to activate complement on gonococci in wild-type mice is likely to be lower than in humans. Ongoing studies have shown efficacy of mAb 2C7 against wild-type strains MS11 and FA1090, which both bind C4BP and when sialylated, also bind FH, in ‘dual’ human FH and C4BP transgenic mice (S.G, P.A.R and S.R, unpublished observations) suggesting that mAb 2C7 can overcome the effects of these complement inhibitors in vivo.

In conclusion, this novel site of sialylation on *N. gonorrhoeae* HepII lactose can inhibit complement activation and also engage Siglecs (45). These findings also explain the ubiquitous expression of HepII lactose (an integral part of the ‘2C7 LOS epitope’) among clinical isolates of *N. gonorrhoeae* and further validate targeting the 2C7 epitope with antibody-based vaccines and immunotherapeutics.

## Materials and Methods

### Bacterial strains

Strain 15253 was recovered from an individual with disseminated gonococcal infection and has been described previously (44). Only *IgtA* and *IgtE* are intact in its lgtA-E locus (69). Deletion of *lgtG* in 15253 to yield 15253/G– has been described previously (70). Deletion of LOS sialyltransferase (lst) to yield 15253 Δlst *(Ist::kan^R^)* was performed as described previously (54). All the LOS mutant derivatives of MS11 have been described previously (48). The LOS phenotypes of 15253, 15253/G– and the MS11 mutants used in this study are listed in Figure 1. Seventy five additional isolates were obtained from subjects enrolled in a transmission study (Ref.) of gonococcal infection from men to women in Nanjing, China. All subjects provided written informed consent in accordance with requirements by Institutional Review Boards from: the University of Massachusetts Medical School; Boston Univeristy School of Medicine and the Institute of Dermatology, Chinese Academy of Medical Sciences & Peking Union Medical College, Nanjing, China.

### Normal human serum

Serum was obtained from normal healthy adult volunteers with no history of gonococcal or meningococcal infection who provided informed consent. Participation was approved by the University of Massachusetts Institutional Review Board for the protection of human subjects. Serum was obtained from whole blood that was clotted at 25 °C for 30 min followed by centrifugation at 1500 g for 20 min at 4 °C. Serum from 10 donors was pooled, aliquoted and stored at −80 °C.

### Mouse complement

Use of animals in this study was performed in strict accordance with the recommendations in the Guide for the Care and Use of Laboratory Animals of the National Institutes of Health. The protocol was approved by the Institutional Animal Care and Use Committee (IACUC) at the University of Massachusetts Medical School. Mouse blood obtained by terminal cardiac puncture was allowed to clot for 20 min at room temperature, then placed on ice for 20 min and centrifuged at 10,000 g for 10 min at 4 °C. Serum was harvested and stored in singleuse aliquots at −80 °C.

### Flow cytometry

Factor H binding to *N. gonorrhoeae* was detected as described previously (71). Briefly, ~10^7^ bacteria in HBSS containing 1 mM CaCl_2_ and 1 mM MgCl_2_ (HBSS++) containing 0.1% BSA was incubated with 10 μg/ml purified human FH (Complement Technologies, Inc.) for 15 min at 37 °C. Bacteria-bound FH was detected with goat anti-human FH (1 μg/ml) (Complement Technologies, Inc.), followed by antigoat IgG FITC (Sigma) at a dilution of 1:100. Bacteria were fixed in 1% paraformaldehyde in PBS and Data were acquired on a FACSCalibur flow cytometer and analyzed using FlowJo software.

Mouse C3 deposition on bacteria was measured by incubating 10^7^ CFU of bacteria in HBSS++/BSA with mouse complement (concentration stated for each experiment) for 20 min at 37 °C. Mouse C3 fragments deposited on bacteria were detected using anti-mouse IgG FITC (MP Biomedicals) at a dilution of 1:100, and flow cytometry was performed as described above.

### SDS-PAGE

LOS in Protease K (Calbiochem)-treated bacterial lysates prepared as described previously (48) in Tricine-SDS Sample Buffer (Boston Biomolecules) was visualized by electrophoresis on Criterion™ 16% Tris-Tricine gels (Bio-Rad) using Tris-Tricine-SDS Cathode buffer (Boston Biomolecules) at 100 V at 4 °C followed by silver staining (Bio-Rad Silver Stain kit).

### Neuraminidase treatment

Desialylation was carried out with α2-3-specific neuraminidase (New England Biolabs; Cat. No. P0743S). Approximately 10^7^ bacteria in GlycoBuffer 1 (New England Biolabs) were treated with 16 U neuraminidase (reaction volume 100 μl) for 1 h at 37 °C. Control reactions contained buffer alone. Bacteria were then incubated with mouse serum as described above to measure quantitatively, C3 deposition.

### Serum bactericidal assay

Strain 15253 was grown in gonococcal liquid media (Morse A, Morse B and IsoVitaleX^TM^ (72)) containing CMP-Neu5Ac at concentrations ranging from 0 to 100 μg/ml in half-log_10_ increments; susceptibility to mAb 2C7 (5 μg/ml) was determined by serum bactericidal assay as described previously (72) with minor modifications. The clinical isolates from Nanjing, China were all grown in liquid media as described above, supplemented with 2 μg/ml of CMP-Neu5Ac. Approximately 2000 CFU gonococci in HBSS_++_/0.1% BSA were incubated with 20% NHS either in the presence or absence of mAb 2C7. Final bactericidal reaction volumes were maintained at 75 μl. Aliquots of 12.5 μl were plated onto chocolate agar plates in duplicate at the beginning of the assay (t_0_) and again after incubation at 37°C for 30 min (t_30_). Survival was calculated as the number of viable colonies at t_30_ relative to t_0_.

### Mass spectroscopic analysis of LOS

O-deacylated LOS was prepared as described previously (26). LC-MS was performed using a Waters Premier Q-TOF operated in the positive-ion mode with an Agilent 1260 capillary LC system. LC separation was done on an Agilent Eclipse XDB C8 column (5μm, 50 × 1mm) operated at 55 ºC. The flow rate was 20 μL/min. Solvent A: aqueous 0.2 % formic acid/0.028 % ammonia; solvent B: Isopropanol with 0.2% formic acid/0.028% ammonia. The following gradient was used: 0-2 min. 10 % B, 2-16 min linear gradient to 85 % B, 16-25 min. 85 %B, 25-30 min. equilibration at 10% B.

### Anti-LOS mAbs

Anti-LOS mAbs 3F11 (73), L8 (74), L1 (75) and 2C7 (42) have been described previously. Fig. 10A indicates the specificities of each of the mAbs

### Whole cell ELISA

Whole cell ELISA was performed as described previously (71). Briefly, U-bottomed microtiter wells (Dynatech Laboratories, Inc., Chantilly, VA) were coated with 50 μl of bacterial suspensions (~10^8^ organisms/ml) in PBS for 3 h at 37 °C, followed by incubation overnight at 4 °C. Plates were washed with PBS containing 0.05% Tween 20. Tissue culture supernatants containing mAbs 3F11, L8, L1 and 2C7 were dispensed into wells and incubated for 1 h at 37 °C, followed by washing with PBS/0.05% Tween 20. Bound 2C7, L8 and L1 were disclosed with anti-mouse IgG alkaline phosphatase (Sigma), and mAb 3F11 was detected with anti-mouse IgM alkaline phosphatase (Sigma).

### Cmah KO mice

Unlike mice, humans lack the ability to convert Neu5Ac to Neu5Gc, because of an *Alu*-mediated deletion in a critical exon that encodes the enzyme, CMP-Neu5Ac hydroxylase (CMAH) (76). Deletion of Cmah in mice results in expression of only Neu5Ac. *Cmah* knockout (KO) mice were generated with a humanlike deletion in exon 6 of Cmah as described previously (77) and were subsequently back-crossed >10 generations into a BALB/c background.

### Mouse infection

The mouse vaginal colonization model developed by Jerse was used (78). Briefly, female *Cmah* KO mice in the diestrus phase of the estrous cycle were started on treatment (that day) with 0.1 mg Premarin^®^ (Pfizer) in 200 μl of water given subcutaneously on each of three days; −2, 0 and +2 days (before, the day of and after inoculation) to prolong the estrus phase of the reproductive cycle and promote susceptibility to *Ng* infection. Antibiotics (vancomycin, colistin, neomycin, trimethoprim and streptomycin (VCNTS)) ineffective against *N. gonorrhoeae* were also used to reduce competitive microflora (7). Mice (n=10/group) were infected on Day 0 with either strain 15253, 15253/G– or 15253 *Δlst* (inoculum specified for each experiment).

Vaginas were swabbed daily and plated on chocolate agar containing VCNTS to enumerate *N. gonorrhoeae* CFUs. The efficacy of mAb 2C7 in vivo against 15253 was performed in wild-type BALB/c mice (Jackson Laboratories) as described previously. Mice were treated with mAb 2C7 or control mouse IgG3 intraperitoneally, 10 μg twice a day on days −2, −1 (prior to) and 0 (the day of infection with strain 15253 and daily vaginal CFU enumeration was carried out as described above.

### Statistical analyses

Experiments that compared clearance of *N. gonorrhoeae* in independent groups of mice estimated and tested three characteristics of the data (43): Time to clearance, longitudinal trends in mean log_10_ CFU and the cumulative CFU as area under the curve (AUC). Statistical analyses were performed using mice that initially yielded bacterial colonies on Days 1 and/or 2. Median time to clearance was estimated using Kaplan-Meier survival curves; times to clearance were compared between groups using the Mantel-Cox log-rank test. Mean log_10_ CFU trends over time were compared between groups using a linear mixed model with mouse as the random effect using both a random intercept and a random slope. A cubic function in time was determined to provide the best fit; random slopes were linear in time. A likelihood ratio test was used to compare nested models (with and without the interaction term of group and time) to test whether the trend differed over time between the two groups. The mean AUC (log_10_ CFU versus time) was computed for each mouse to estimate the bacterial burden over time (cumulative infection); the means under the curves were compared between groups using the nonparametric two-sample Wilcoxon rank-sum (Mann-Whitney) test because distributions were skewed or kurtotic. The Kruskal-Wallis equality-of-populations rank test was also applied to compare more than two groups in an experiment. Correlation between survival in serum bactericidal assays and mAb 2C7 binding was performed by Spearman’s non-parametric test. A cubic equation as used to generate the best-fit curve.

## Acknowledgments

We thank Nancy Nowak and Samuel Fountain for technical assistance. This work was supported by grants from the National Institutes of Health / National Institutes of Allergy and Infectious Diseases, AI118161 (to S.R.), AI114790, AI132296 (to P.A.R and S.R.) and AI114710 (to P.A.R. and S.G.).

